# Identifying The “Core” Transcriptome of SARS-CoV-2 Infected Cells

**DOI:** 10.1101/2021.09.22.461142

**Authors:** Elanood Tageldin Nour, Ryan Tran, Ayda Afravi, Xinyue Pei, Angela Davidian, Pavan Kadandale

## Abstract

In 2019, the severe acute respiratory syndrome coronavirus 2 (SARS-CoV-2) first emerged, causing the COVID-19 pandemic. Consequently, ongoing research has focused on better understanding the mechanisms underlying the symptoms of this disease. Although COVID-19 symptoms span a range of organ systems, the specific changes in gene regulation that lead to the variety of symptoms are still unclear. In our study, we used publicly available transcriptome data from previous studies on SARS-CoV-2 to identify commonly regulated genes across cardiomyocytes, human bronchial epithelial cells, alveolar type II cells, lung adenocarcinoma, human embryonic kidney cells, and patient samples. Additionally, using this common “core” transcriptome, we could identify the genes that were specifically and uniquely regulated in bronchial epithelial cells, embryonic kidney cells, or cardiomyocytes. For example, we found that genes related to cell metabolism were uniquely upregulated in kidney cells, providing us with the first mechanistic clue about specifically how kidney cells may be affected by SARS-CoV-2. Overall, our results uncover connections between the differential gene regulation in various cell types in response to the SARS-CoV-2 infection and help identify targets of potential therapeutics.

## Introduction

COVID-19 is a contagious disease caused by severe acute respiratory syndrome coronavirus 2 (SARS-CoV-2), a positive sense, single-stranded RNA virus. Since 2019, COVID-19 has caused millions of deaths globally and has led to devastating effects on human society both economically and socially. The development of effective vaccines has greatly mitigated the spread of COVID-19. However, there has been a continuous spread of misinformation regarding these vaccines, as well as a significant portion of the population that refuses vaccination. As a result, new cases continue to be reported across the world, which has led to the rise of new, more infective variants of the virus, hampering efforts to control the disease [1]. For example, the exponential increase in positive cases in India in April 2021 coincided with the prevalence of new SARS-CoV-2 variants [2]. Thus, despite the availability of potent vaccines, the biological and sociopolitical factors have prevented us from eliminating the threat of SARS-CoV-2 transmission and necessitated a need for the development of effective treatments to help reduce the severity of this disease. SARS-CoV-2 is characterized by rapid human-to-human transmission from droplet contamination [3, 4, 5]. The viral infection causes a variety of symptoms ranging from mild to severe in different patients. These symptoms are due to the virus affecting a number of different organs within the body, including the pulmonary, cardiovascular and renal systems. Some of the symptoms even persisted after recovery from the viral infection [6, 7, 8]. Thus, identification of the mechanisms by which SARS-CoV-2 affects different organs will provide us with targets for the development of effective therapeutic strategies, synergizing with the ongoing effort to eliminate the virus by mass vaccinations.

In order to gain access to the host cell, SARS-CoV-2 binds to the Angiotensin Converting Enzyme 2 (ACE2) receptor, which is highly expressed in the lower respiratory tract, and proceeds with viral replication [9]. After entering the cell by endocytosis, the virus and the host cell continue to interact intensively. The virus utilizes host cell machinery to transcribe its genome, and the host pattern recognition receptors (PRR) also recognize pathogen-associated molecular patterns (PAMP) derived from viral proteins or genome, triggering the innate immune response [10]. For example, in lung cells, viral infection increases the secretion of pro-inflammatory cytokines such as IL-1β and IL-6 [11, 12].

Aside from epithelial tissue in the respiratory tract, other tissues also express ACE2 receptors, such as intestine, kidney, stomach, colon, and heart muscle, suggesting their potential to be infected and lead to tissue-specific symptoms, such as diarrhea and vomiting resulting from gastrointestinal dysfunction [13]. Depending on the type of cell, the post-infection phenotype of different cells may vary. For example, SARS-CoV-2 infected neurons showed decreased expression of brain-derived neurotrophic factor (BDNF), which is important in neuronal connection [14], while human iPSC-derived cardiomyocytes cease beating 72 hours after infection [15]. The different responses of different cell types complicate the ability to study the comprehensive effects of SARS-CoV-2 in the human body or to develop effective therapeutics against the disease.

Recent work has uncovered the unique properties of the immune response to SARS-CoV-2 [16]. Other studies have uncovered how SARS-CoV-2 changes the transcriptome of various cell types, including Caco-2 [17], Vero, Huh-7, 293T, A549, and Efk3B [18]. However, no one, to date, has compared differential gene expression across cell types from different organ systems to uncover the “core” changes to gene expression caused by viral infection across all cell types. Observing differential gene expression in cell lines provides insight into the unique interactions and effects of the virus on the transcriptome of specific/individual cells. By combining these datasets, we can address what genes are commonly regulated due to viral infection across cell types, establishing a “core” viral transcriptome. Comparing the differentially regulated genes in a specific cell type to this “core” transcriptome will uncover the genes that are uniquely regulated in that cell type, yielding insights into the specific effects of the virus on these cells. In this study, using publicly available transcriptome data from previous studies on SARS-CoV-2 involving multiple cell lines and patient samples, we obtained targets of the virus that are consistent among multiple cell types. This helps to provide a model for multiple (rather than a single) body systems/organs and their responses to COVID-19 [19]. Using these findings, we can examine cell-intrinsic and cell-extrinsic responses to infection by SARS-CoV-2 [19] and identify clinical risk factors.

## Materials and Methods

### Choosing BioProjects

Human transcriptomic data for uninfected and SARS-CoV-2 infected samples were obtained from the NCBI BioProject database. Search results were filtered for SRA (Sequence Read Archive) data derived from Illumina sequencing. We chose to examine data from only a single RNA sequencing methodology in order to filter out any biases that may be intrinsic to the method used. A BioProject was chosen for the study if there were at least two replicates (at least two different SRR numbers) for both uninfected and SARS-CoV-2 infected samples. The type of sample (type of cell line or patient sample), whether the sample was single-end or paired-end, and time after infection were noted for each SRR.

### Galaxy workflow

The Galaxy platform was used to carry-out the analysis [20]. The key steps in the Galaxy workflow are FASTQC, Trimmomatic, HISAT2, and FeatureCounts. First, FASTQC is run on the datasets to check for the sequence files’ qualities. Second, the Trimmomatic tool is used to remove Illumina adapter sequences from the reads, to trim the low-quality sequences from either end of the reads, and to remove any sequences with a less than 25 nt length. Third, HISAT2 gives the overall alignment rate, thereby allowing the user to know how much of each sequence file maps back to the human genome. Finally, FeatureCounts generates counts tables for the sequence files. A one-factor DESeq2 analysis is then performed on each of the samples separately to examine the replicates and exclude the low quality datasets (see below).

### Choosing SRRs

All SRRs present in a particular BioProject for both uninfected and SARS-CoV-2 infected samples were selected for an initial run-through of the experimental workflow in Galaxy. A PCA graph generated by a one-factor DESeq2 analysis determined whether there was a clear separation between uninfected and infected samples in order to demonstrate that there was a distinct difference between uninfected and infected samples. Samples that did not show separation were subsequently removed from the study and excluded from further analyses. The SRRs selected for this study (Table S1) were from patient lung samples and eight cell lines: human embryonic kidney 293 cells with SV40 large T antigen (HEK 293T), human induced pluripotent stem cell-derived cardiomyocytes (hiPSC-CMs), human induced pluripotent stem cell-derived alveolar type II epithelial-like cells (iAT2s), iPSCs, Vero E6, primary human bronchial epithelial cells (NHBE), human alveolar epithelial cells (lung adenocarcinoma) located on basal side (A549), and human bronchial epithelial cells (lung adenocarcinoma) located on apical side (Calu-3).

### Combined analysis

After obtaining FeatureCounts tables for all replicates of all samples, we conducted a two-factor DESeq2 analysis (combined analysis) across all eight cell lines and patient lung samples in order to identify commonly differentially expressed genes upon viral infection. The two factors used were “*Infection*” and “*Sample Type*”. The first factor, “*Infection*”, had two levels: “*Uninfected*” and “*Infected*”. This allowed us to find the differences in gene expression between the uninfected and infected samples (regardless of sample type). The second factor, “*Type*”, had nine levels, corresponding to the different samples that we were analyzing. This accounted for the inherent differences in gene expression between all samples.

### Identifying differentially regulated genes that are unique to the cell lines

To identify differentially regulated genes that are unique to the cell lines, the DESeq2 results from the combined analysis were compared with those from the cell line analyses. First, a one-factor DESeq2 analysis was performed on each three selected non-cancerous cell lines: NHBE, HEK 293T, and hiPSC-CM. In each DESeq2 analysis, the counts tables (generated from the FeatureCounts step) of the replicates of a cell line were compared based on one factor, “*Infection*”, with two levels: “*Uninfected*” and “*Infected*”. The goal of this factor was to uncover the differences in gene expression between the uninfected and infected samples of a cell line. In order to screen for differentially regulated genes that are unique to each cell line, the upregulated and downregulated genes from each of the three selected cell lines were compared with the corresponding upregulated and downregulated genes from the combined analysis using Microsoft Excel.

### Determining biological significance of the “core” transcriptome using g:Profiler and R code to generate PCA plots and clusters of GO terms

Given the too small sample size, the uniquely downregulated genes from the HEK 293T cell line analysis as well as both the upregulated and downregulated genes from the NHBE cell line analysis were examined individually in order to understand the likely biological role of these genes in viral infection. On the other hand, given the sheer number of differentially expressed genes from the combined analysis (upregulated genes), HEK 293T cell line analysis (uniquely upregulated genes), and the hiPSC-CM cell line analysis (both uniquely upregulated and downregulated genes), it was particularly laborious to individually analyze each gene. Instead, a gene ontology (GO) analysis was performed on those sets of results by inputting each list of genes into g:Profiler, a web server for functional enrichment analysis, to obtain gene ontology (GO) terms from each of the three sub-ontologies: Biological Process (BP), Cellular Component (CC), and Molecular Function (MF) [21]. The results returned an overwhelming number of GO terms, which were then analyzed using R [22], following the procedure described below.

Each set of differentially regulated genes that could not be examined and analyzed individually was put through a pipeline that produced clusters of GO terms. First, the list of gene IDs from a set was input into g:Profiler in order to find the GO terms associated with each differentially regulated gene. g:Profiler matches the input gene IDs to GO terms from each of the three GO term categories or sub-ontologies: Biological Process (BP), Cellular Component (CC), and Molecular Function (MF). We then filtered the GO terms for those that were significantly overrepresented in each list of genes (P_adj_ < 0.01). In order to find what biological functions, in general, were represented by the differentially regulated genes, we converted the list of GO terms into a binary matrix in which the columns were the GO terms, and each row (corresponding to a gene) was a binary set of numbers that represent whether a given GO term was associated with that gene (represented by “1” in the binary matrix) or not (“0” in the binary matrix). This binary matrix was then used to perform hierarchical clustering in R [22], to uncover genes with likely similar functions (based on their GO terms being closely grouped together by the hierarchical clustering).

To visualize the clustering results, we created a PCA plot to examine whether the results of the clustering were reasonable, and to determine the best number of clusters to use for a given dataset. To achieve this, we aimed to obtain PCA plots that exhibited the clearest separation between the clusters of genes while minimizing the variance between the points within a cluster. When a cluster comprised more than 75 genes, we sub-clustered this set of genes, using the same method described above.

## Results

### Quality control of datasets

To test the quality of the individual datasets collected for the purposes of this experiment, a DESeq2 analysis was done on each dataset. Using Principal Component Analysis (PCA) plots (Figures 1 and 2) from the DESeq2 data, the quality of datasets were determined by examining the pattern of clustering for the data points (each data point corresponds to an SRR dataset).

**Figure 1.**
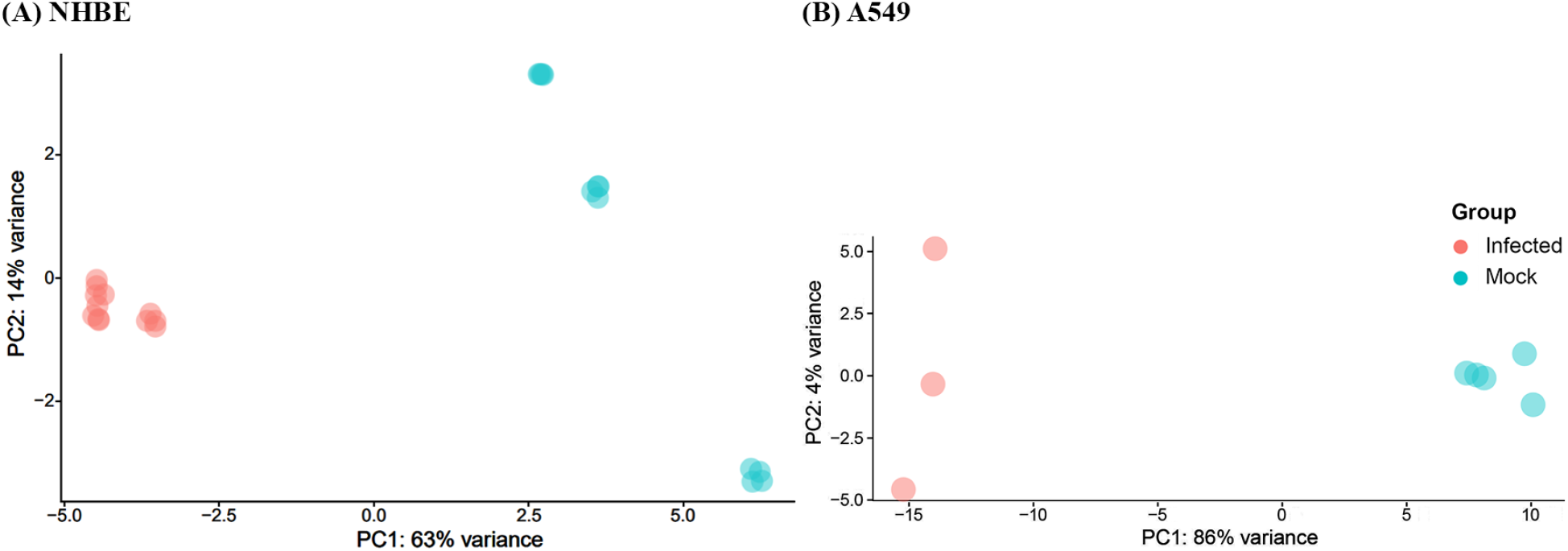
Principal Component Analysis (PCA) Plots for exemplar high quality datasets. All plots show high variance between mock vs. infected datasets. The figures are PCA plots for NHBE **(A)** and A549 **(B)** datasets. There is high variance between mock and infected datasets.

**Figure 2.**
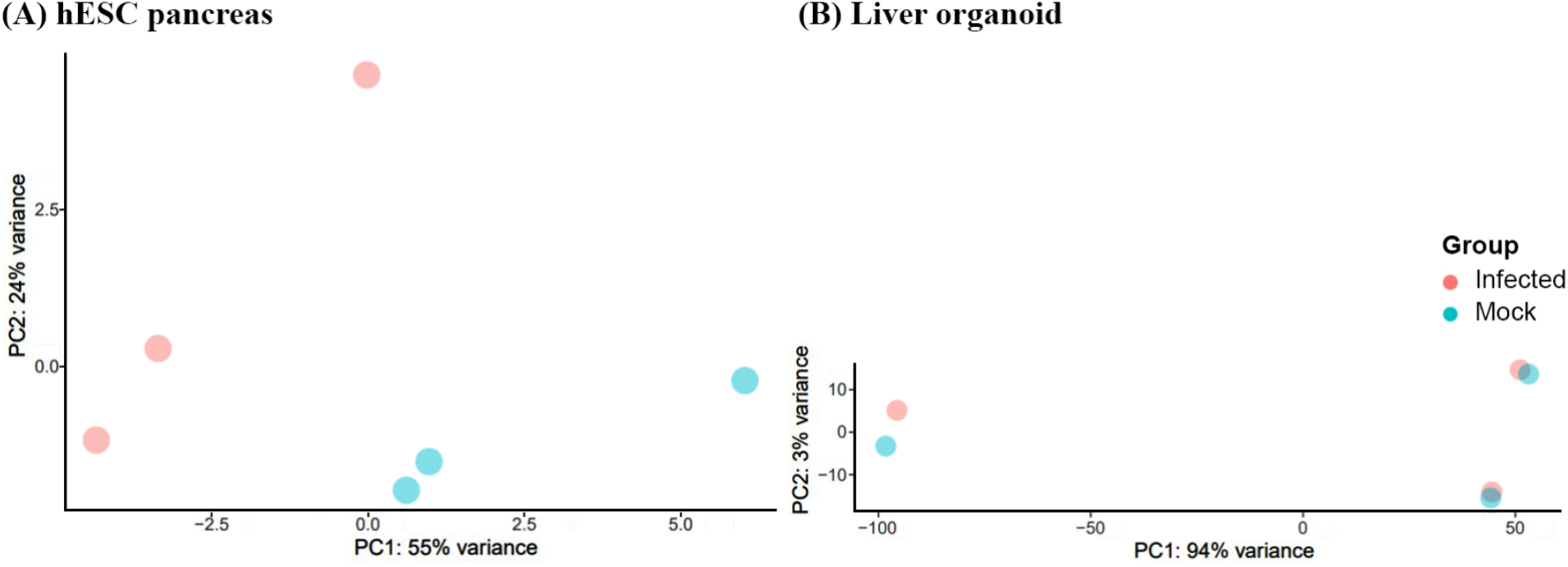
Excluded Principal Component Analysis (PCA) Plots. All plots included in this figure proved to be outliers to the collective data collected in the experiment. (A) hESC mock and infected datasets had insignificant variance and almost no correlation within similar factors. (B) Liver Organoid plot had no clear separation between mock and infected datasets.

High quality datasets were those in which the infected samples clustered together, and away from the mock (or un-) infected samples—for example, the NHBE (Figure 1A) and A549 cell lines (Figure 1B). Finally, using the same quality measures, we rejected the low-quality datasets, including a hESC pancreas cell line (Figure 2A) and liver organoid (Figure 2B), in which there was no clear separation between the infected and uninfected samples. This quality control step ensures that any variance in the final results will be primarily due to changes caused by the virus, rather than discrepancies within the datasets themselves.

After passing the quality control step, all individual samples were compiled to perform a single combined DESeq2 analysis for the purpose of performing a comparative RNA-seq analysis across all samples.

### Identification of the “core” (commonly up and downregulated genes) transcriptome

We combined the high quality datasets and performed a two-factor DESeq2 analysis in order to identify genes that were commonly upregulated or downregulated across all cell types. The PCA plot from the combined data (Figure 3) showed that cell type mattered more than infection status, indicating that differential gene regulation upon viral infection did not supersede the similarities in gene expression within each cell type. From this DESeq2 analysis, we identified commonly up and downregulated genes across all cell types as those that had a fold-expression change greater than 2 (or less than −2) - indicated by the black lines in Figure 4, and which had an adjusted P-value less than 0.05 (Figure 4, red dots). Based on these criteria, we found 866 genes that were significantly differentially upregulated in the SARS-CoV-2 infected samples, while there are only 9 genes that were significantly differentially downregulated in the infected samples (Table S2).

**Figure 3.**
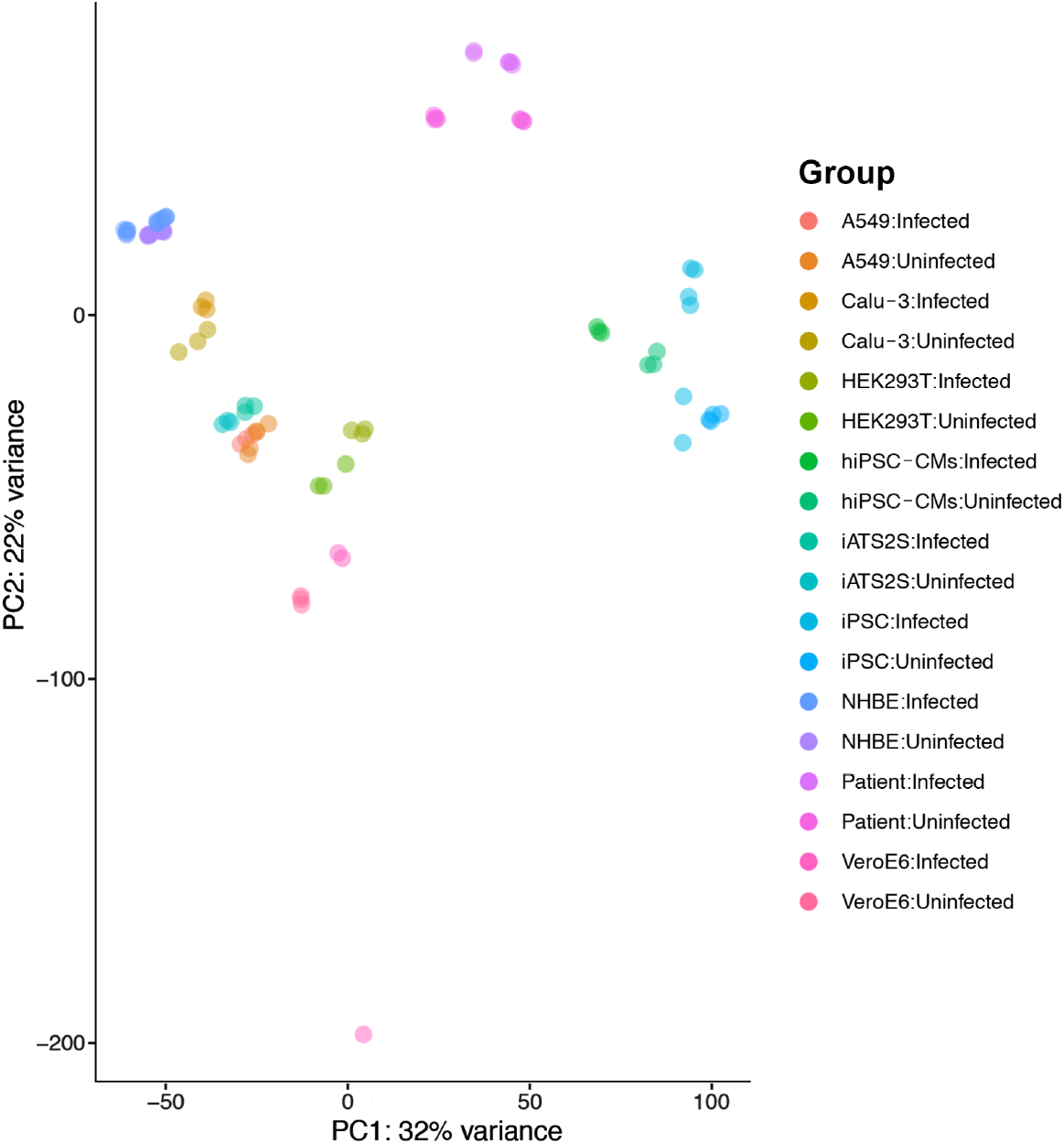
Combined Principal Component Analysis (PCA) Plot. Cell types and patient samples showed distinct grouping, indicating type factor dominates over infection factor.

**Figure 4.**
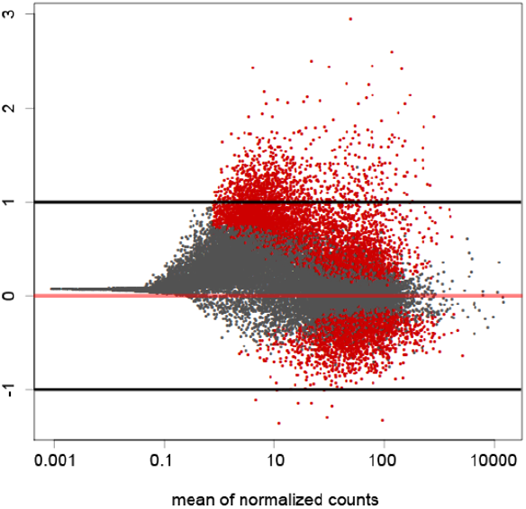
Combined MA Plot. This figure plots the differentially regulated genes according to their Log_2_ Fold Change (Log_2_FC) values. The Red dots represent the genes that were significantly differentially up- or downregulated, while the Grey dots represent the genes that were similarly expressed and had an approximate Log_2_FC value closer to 0.

In order to understand the biological significance of these upregulated genes, we used g:Profiler to find GO term annotations that were significantly enriched (P_adj_ < 0.01) in this set of 866 genes. From this, we obtained a total of 199 total GO terms (Table S3A), of which 169 were Biological Process (BP) terms, 13 were Cellular Component (CC) terms, and 17 were Molecular Function (MF) terms) corresponding to the known functions of these genes. Hierarchical clustering followed by PCA analysis resulted in these 199 terms forming 6 major clusters (Figure 5 and Table S4A). These clusters were, broadly, “Immune, defense, and inflammatory response”, “Leukocyte activation and secretion”, “Cytokine activity and response”, “Leukocyte activation and cell adhesion”, “Response to stimulus, signaling, and communication”, and “Ion transporter activity” (Figure 5).

**Figure 5.**
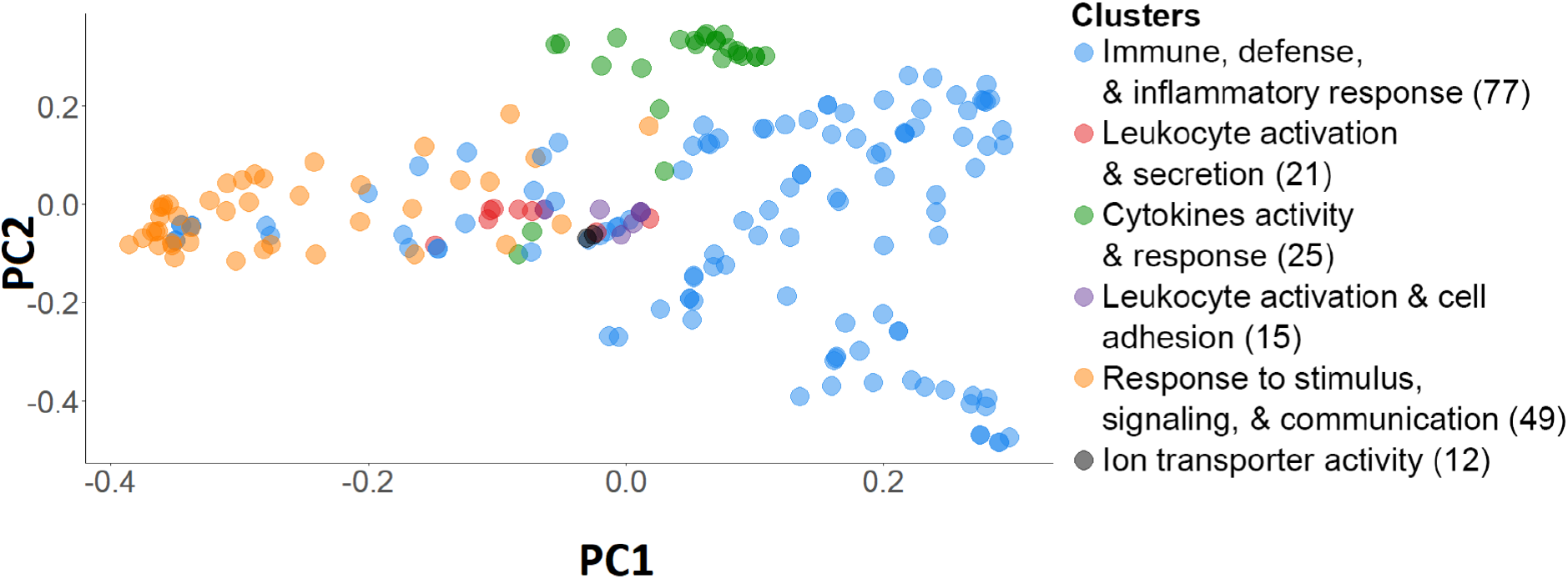
Combined Principal Component Analysis (PCA) plot for 199 GO terms (P_adj_ < 0.01) of the commonly upregulated genes. There were 866 upregulated genes, 773 of which were recognized by g:Profiler. They are categorized into six clusters, represented with different colors, using the R code with Jaccard algorithm. The number of GO terms within the clusters is displayed next to each cluster.

Cluster 1 (“Immune, defense, and inflammatory response”) contained over 75 GO terms (77) and was further split into 8 sub-clusters (Figure S2 and Table S4B). The resulting sub-clusters were “Interleukin-6 and TNF production and regulation”, “Interleukin-10 and interferon-gamma production and leukocyte differentiation and activation”, “Inflammatory and defense response”, “Response to virus”, “Regulation of immune response”, “Acute inflammatory response and receptor signaling”, “Cytokine production”, and “Leukocyte migration and response to interleukin-1 and interferon-gamma” (Table 1). 5 of the 6 GO term clusters, including the 8 sub-clusters, are related to immune response. The cluster that is not involved in immune response is “Ion transporter activity”.

**Table 1.**
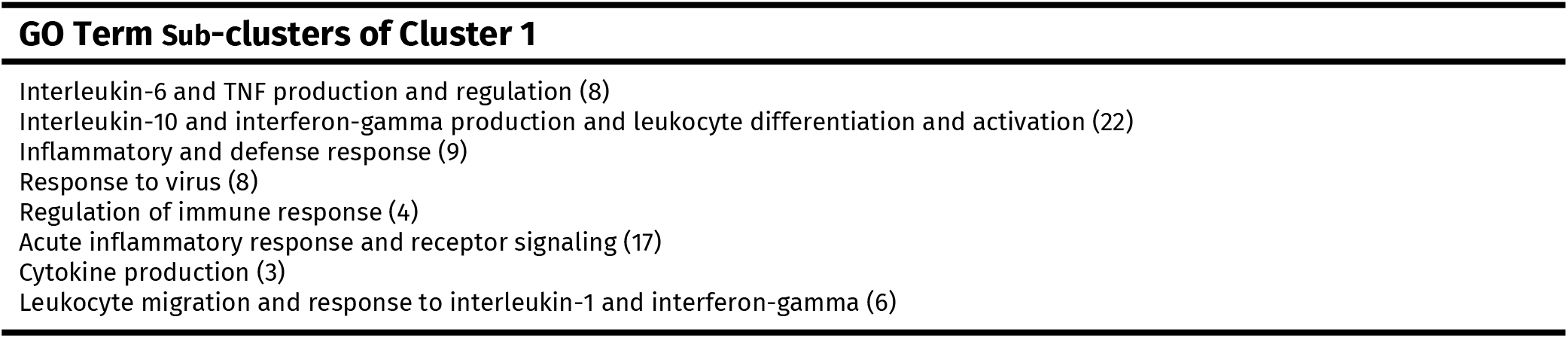
GO term sub-clusters of Cluster 1 from the upregulated results of the combined analysis. The names of the eight GO term sub-clusters from the upregulated results of the combined analysis are displayed in the table. The number of GO terms that are in each sub-cluster is in parentheses.

On the other hand, only nine commonly downregulated genes were identified by the two-factor DESeq2 analysis across all samples (Table S2B). Therefore, rather than using g:Profiler to find GO term annotations as from the analysis of the upregulated results, the nine commonly downregulated genes were examined individually in order to understand the biological significance of these genes in viral infection (Table 2). The nine genes spanned a wide range of biological functions, including inhibition of endothelial permeability, contractile activity, cellular growth, organization of cytoskeleton, and cell differentiation. For example, NME5 is responsible for protecting the cell from apoptosis by Bax, and Retinoic acid-induced protein 2 (RAI2) is involved in Type-I IFN synthesis and release pathways [23, 24].

**Table 2.** Downregulated genes from the combined analysis. Nine genes were commonly downregulated across all cell lines and patient samples. The genes were obtained by filtering the two-factor DESeq2 analysis results for significance and downregulation (P_adj_ < 0.01; Log_2_FC < −1). The GO terms were obtained from g:Profiler, using a P-value threshold of 1.00 to retrieve the terms for a single gene.

| Gene ID | Gene Name | GO Terms |
| --- | --- | --- |
| 284 | Angiopoietin-1 (ANGPT1) | MF: Protein tyrosine kinase binding<br>BP: Regulation of macrophage migration inhibitory factor signaling pathway;<br>Positive regulation of blood-brain barrier permeability<br>CC: Extracellular organelle |
| 4638 | Myosin light chain kinase (MYLK) | MF: Myosin light chain kinase activity<br>BP: Aorta smooth muscle tissue morphogenesis; Cellular hypotonic response<br>CC: Cleavage furrow |
| 8382 | NME5 | MF: Nucleoside diphosphate kinase activity<br>BP: UTP biosynthetic process; GTP biosynthetic process<br>CC: Extracellular region |
| 9452 | Integral membrane protein 2A (ITM2A) | MF: Amyloid-beta binding<br>BP: Plasma cell differentiation<br>CC: Cell periphery |
| 10742 | Retinoic acid-induced protein 2 (RAI2) | MF: Binding<br>BP: Multicellular organism development<br>CC: Cellular component |
| 11013 | Thymosin beta 15A (TMSB15A) | MF: Actin monomer binding<br>BP: Sequestering of actin monomers<br>CC: Intracellular non-membrane-bounded organelle |
| 25840 | Methyltransferase like 7A (METTL7A) | MF: Catalytic activity<br>BP: Cell activation involved in immune response<br>CC: Tertiary granule lumen |
| 283481 | FGF14 antisense RNA 2 (FGF14-AS2) | MF: N/A<br>BP: N/A<br>CC: N/A |
| 729967 | MORN repeat containing 2 (MORN2) | MF: Binding<br>BP: Developmental process involved in reproduction; Spermatogenesis<br>CC: Secretory vesicle |

### Differentially regulated genes in specific cell types

Having identified the core transcriptome of commonly up and downregulated genes, we sough to identify genes that were differentially regulated in different cell types that were unique to those cells. This will help us parse out the specific effects of SARS-CoV-2 on different cell types and organ systems. As proof of principle of our approach we chose a lung-derived, a kidney-derived, and cardiac-derived cell type, and uncovered genes that were uniquely up and downregulated by viral infection in these cells.

In order to compare the gene expression in different cell types (bronchial epithelial cells, embryonic kidney cells, and cardiomyocytes), a single-factor DESeq2 analysis was each performed on the NHBE, HEK 293T, and hiPSC-CM cell lines, respectively, and the resulting differentially regulated genes of each cell line were compared with the commonly regulated genes to obtain the uniquely regulated genes for each cell type (Table 3).

**Table 3.** Number of differentially regulated genes that are unique to the NHBE, HEK 293T, and hiPSC-CM cell lines. The differentially regulated genes for each cell line were obtained by performing the DESeq2 analysis in Galaxy (P_adj <_ 0.01 and Log_2_FC < −1 or Log_2_FC > 1). The table shows the total number of up- and down-regulated genes identified, and the number of unique genes (that do not show up in the common set of up- or down-regulated genes from the combined analysis) for each cell type analyzed.

|  | Total no. of genes |  | No. of unique genes |  | % of genes unique to cell line |  |
| --- | --- | --- | --- | --- | --- | --- |
|  | UP | DOWN | UP | DOWN | UP | DOWN |
| NHBE | 60 | 4 | 12 | 3 | 20.0% | 75.0% |
| HEK 293T | 321 | 12 | 262 | 12 | 81.6% | 100% |
| hiPSC-CM | 1482 | 1860 | 1257 | 1854 | 84.8% | 99.7% |

For the uniquely downregulated genes from the HEK 293T cell line, as well as both the upregulated and downregulated genes from the NHBE cell line we examined the genes individually, since the number of genes in each case was fairly small. In order to analyze the large number of hits for the uniquely upregulated genes from the HEK 293T cell line as well as both the upregulated and downregulated genes from the hiPSC-CM cell line, we used GO term clustering as described previously, in order to understand the biological relevance of these genes being differentially regulated upon viral infection.

### Genes that are differentially regulated specifically in NHBE cells

For the NHBE cell line, 12 genes were uniquely upregulated, while 3 genes were uniquely downregulated (Table S5A). Among the 12 upregulated genes (Table 4), 5 genes have functions that are related to immune responses. These 5 genes are interleukin 32 (IL32), LIF interleukin 6 family cytokine (LIF), 2′-5′-oligoadenylate synthetase 3 (OAS3), serpin family A member 3 (SERPINA3), and complement C3 (C3). Other genes, such as hephaestin like 1 (HEPHL1), solute carrier family 6 member 14 (SLC6A14), and Rh family C glycoprotein (RHCG), are involved in transporters activities. 4 genes, MAF bZIP transcription factor F (MAFF), heparin binding EGF like growth factor (HBEGF), TNFAIP3 interacting protein 1 (TNIP1), plasminogen activator tissue type (PLAT), participate in multiple cell signaling activities. Of the 3 downregulated genes (Table 5), C-X-C motif chemokine ligand 14 (CXCL14) takes a part in immune response, while the other 2 genes, interferon induced transmembrane protein 10 (IFITM10) and MAP7 domain containing 2 (MAP7D2), have not been widely studied by the scientific community.

**Table 4.** Uniquely upregulated genes for NHBE. 12 unique genes are differentially upregulated from the NHBE cell line analysis. The genes were obtained by performing the DESeq2 in Galaxy (P_adj_ < 0.01; Log_2_FC > 1). The Gene Ontology terms were obtained from g:Profiler, using a P-value threshold of 1.00 to retrieve the terms for a single gene.

| Gene ID | Gene Name | GO Terms |
| --- | --- | --- |
| 23764 | MAF bZIP transcription factor F (MAFF) | MF: Transcription cis-regulatory region binding<br>BP: Parturition; Chordate embryonic development<br>CC: Chromatin |
| 11254 | solute carrier family 6 member 14 (SLC6A14) | MF: Alanine transmembrane transporter activity<br>BP: Alanine transport<br>CC: Extracellular vesicle |
| 341208 | hephaestin like 1 (HEPHL1) | MF: Ferroxidase activity<br>BP: Copper ion transport<br>CC: Plasma membrane |
| 9235 | interleukin 32 (IL32) | MF: Receptor ligand activity<br>BP: Positive regulation of type III interferon production<br>CC: Cellular anatomical entity |
| 10318 | TNFAIP3 interacting protein 1 (TNIP1) | MF: Mitogen-activated protein kinase binding<br>BP: Modulation by symbiont of host I-kappaB kinase/NF-kappaB cascade<br>CC: Intracellular organelle lumen |
| 3976 | LIF interleukin 6 family cytokine (LIF) | MF: Leukemia inhibitory factor receptor binding<br>BP: Positive regulation of histone H3-K27 acetylation<br>CC: Cellular anatomical entity |
| 1839 | heparin binding EGF like growth factor (HBEGF) | MF: Epidermal growth factor receptor binding<br>BP: Regulation of elastin biosynthetic process<br>CC: Clathrin-coated endocytic vesicle membrane |
| 4940 | 2'-5'-oligoadenylate synthetase 3 (OAS3) | MF: 2'-5'-oligoadenylate synthetase activity; Adenylyltransferase activity<br>BP: Chemokine (C-X-C motif) ligand 9 production<br>CC: Intracellular organelle lumen |
| 5327 | plasminogen activator, tissue type (PLAT) | MF: Catalytic activity<br>BP: Trans-synaptic signaling by BDNF, modulating synaptic transmission<br>CC: Cell junction; Secretory granule |
| 51458 | Rh family C glycoprotein (RHCG) | MF: Ammonium transmembrane transporter activity; Ankyrin binding<br>BP: Transepithelial ammonium transport<br>CC: Extracellular vesicle |
| 12 | serpin family A member 3 (SERPINA3) | MF: Nucleic acid binding<br>BP: Regulation of proteolysis<br>CC: Platelet alpha granule lumen |
| 718 | complement C3 (C3) | MF: C5a anaphylatoxin chemotactic receptor binding<br>BP: Positive regulation of type II hypersensitivity<br>CC: Extracellular vesicle |

**Table 5.** Uniquely downregulated genes for NHBE. Three unique genes are differentially downregulated from the NHBE cell line analysis. The genes were obtained by performing the DESeq2 in Galaxy (P_adj_ < 0.01; Log_2_FC < −1). The Gene Ontology terms were obtained from g:Profiler, using a P-value threshold of 1.00 to retrieve the terms for a single gene.

| Gene ID | Gene Name | GO Terms |
| --- | --- | --- |
| 402778 | interferon induced transmembrane protein 10 (IFITM10) | MF: N/A<br>BP: Plasma membrane<br>CC: N/A |
| 9547 | C-X-C motif chemokine ligand 14 (CXCL14) | MF: G protein-coupled receptor binding<br>BP: Positive regulation of natural killer cell chemotaxis<br>CC: Intracellular membrane-bounded organelle |
| 256714 | MAP7 domain containing 2 (MAP7D2) | MF: N/A<br>BP: Microtubule-based process<br>CC: Intracellular organelle |

### Genes that are differentially regulated specifically in HEK 293T cells

The 262 uniquely upregulated genes in HEK 293T cells (Table S5B) resulted in 62 total GO term annotations (37 BP terms, 3 CC terms, and 22 MF terms) that were significantly enriched in this list (Table S3B). This set of GO terms formed 5 clusters, which were broadly defined as “Transcription and DNA binding”, “RNA biosynthetic process”, “Ion binding”, “Dynein and transcription repressor activity”, and “Cyclic compound binding” (Table 6, Figure S3, and Table S6). All 5 clusters had GO annotations unique to this cell type, and were not seen in the set of genes upregulated across all cell types, showing that these functions were being uniquely regulated in these cells.

**Table 6.**
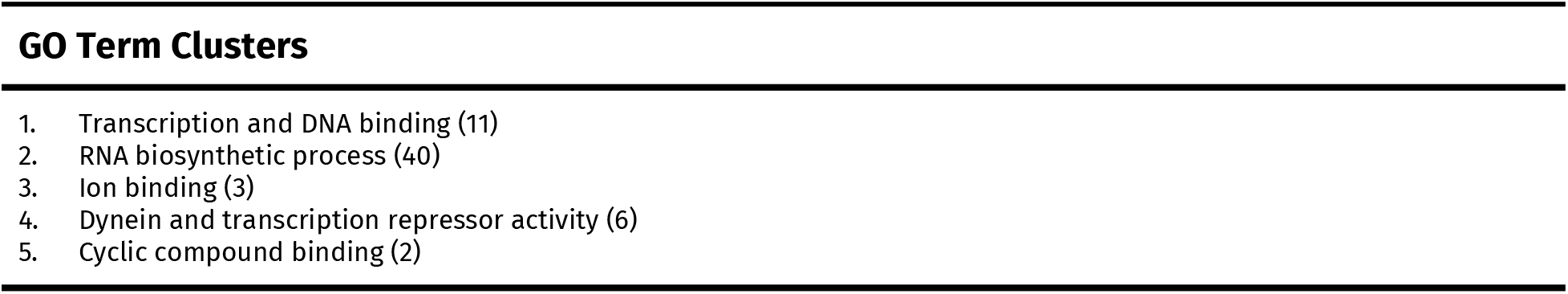
GO term clusters from the upregulated results of the HEK 293T cell line analysis. There were 262 uniquely upregulated genes from the HEK 293T cell line analysis, of which 253 were recognized by g:Profiler. The names of the five GO term clusters are displayed in the table. The number of GO terms that are in each cluster is in parentheses.

Compared to the 9 commonly downregulated genes across all cell lines and patient samples, 12 genes were uniquely downregulated in the HEK 293T cell line (Tables 7 and S5B). Among the 12 genes, 2 genes have not been characterized and 2 genes do not have corresponding gene ontology terms as their functions might not be fully studied. 3 genes, fucose-1-phosphate guanylyltransferase (FPGT), pyruvate dehydrogenase kinase 1 (PDK1), and metabolism of cobalamin associated A (MMAA), play a role in metabolic processes. Solute carrier family 39 member 10 (SLC39A10), which is related to zinc ion transporter activities, and STEAP family member 1 (STEAP1), which participates in ion metabolism, were also revealed to be downregulated. The rest of genes, progestin and adipoQ receptor family member 8 (PAQR8), BCL2 interacting protein 3 like (BNIP3L), and UDP glycosyltransferase 8 (UGT8), all take part in distinct cellular processes, including hormone receptor activities, mitochondrial apoptosis, and galactosyltransferase activities.

**Table 7.** Uniquely downregulated genes for HEK 293T. 12 unique genes are differentially downregulated from the HEK 293T cell line analysis. The genes were obtained by performing the DESeq2 in Galaxy (P_adj_ < 0.01; Log_2_FC < −1). The Gene Ontology terms were obtained from g:Profiler, using a P-value threshold of 1.00 to retrieve the terms for a single gene.

| Gene ID | Gene Name | GO Terms |
| --- | --- | --- |
| 85315 | progesterone and adiponectin receptor family member 8 (PAQR8) | MF: Steroid hormone receptor activity<br>BP: Response to chemical<br>CC: Intracellular anatomical structure |
| 8790 | fucose-1-phosphate guanylyltransferase (FPGT) | MF: Fucose-1-phosphate guanylyltransferase activity<br>BP: Fucose metabolic process<br>CC: Cytosol |
| 57181 | solute carrier family 39 member 10 (SLC39A10) | MF: Zinc ion transmembrane transporter activity<br>BP: Positive regulation of protein tyrosine phosphatase activity<br>CC: Plasma membrane |
| 257396 | novel transcript, antisense to MOCS2 (nan) | MF: N/A<br>BP: N/A<br>CC: N/A |
| 26872 | STEAP family member 1 (STEAP1) | MF: Catalytic activity; Transporter activity<br>BP: Transport<br>CC: Membrane-bounded organelle |
| 5163 | pyruvate dehydrogenase kinase 1 (PDK1) | MF: Pyruvate dehydrogenase (acetyl-transferring) kinase activity<br>BP: Hypoxia-inducible factor-1alpha signaling pathway; Regulation of acetyl-CoA biosynthetic process from pyruvate<br>CC: Mitochondrial pyruvate dehydrogenase complex |
| 105379045 | nan (nan) | MF: N/A<br>BP: N/A<br>CC: N/A |
| 79940 | long intergenic non-protein coding RNA 472 (LINC00472) | MF: N/A<br>BP: N/A<br>CC: N/A |
| 79896 | threonine synthase like 1 (THNSL1) | MF: N/A<br>BP: N/A<br>CC: N/A |
| 166785 | metabolism of cobalamin associated A (MMAA) | MF: Nucleotide binding<br>BP: Cobalamin biosynthetic process; Short-chain fatty acid catabolic process<br>CC: Intracellular organelle lumen |
| 7368 | UDP glycosyltransferase 8 (UGT8) | MF: N-acylsphingosine galactosyltransferase activity<br>BP: Protein localization to paranode region of axon; Paranodal junction assembly<br>CC: Cell periphery |
| 665 | BCL2 interacting protein 3 like (BNIP3L) | MF: Lamin binding<br>BP: Mitochondrial protein catabolic process<br>CC: Intrinsic component of membrane |

### Genes that are differentially regulated specifically in hiPSC-CM cells

We found 1482 upregulated genes from the hiPSC-CM cell line, of which 1257 were unique (Table S5C). These resulted in 422 total GO terms (386 BP terms, 16 CC terms, and 20 MF terms) (Table S3C) that could be collapsed into 8 clusters (Table 8, Figure S4, and Table S7A). 4 of these clusters were also found in the commonly upregulated genes: “Immune response, transcription, and oxidoreductase activity”, “Regulation of signaling, communication, cell death, protein metabolic process, and catalytic activity”, “Response to stimulus, signaling, and communication”, and “Immune, defense, and inflammatory response”. The other 4 GO term clusters were unique to these cells: “Regulation of transcription and biosynthetic processes”, “Binding and metabolic process”, “Angiogenesis and cell motility”, and “Regulation of cellular metabolic process”.

**Table 8.**
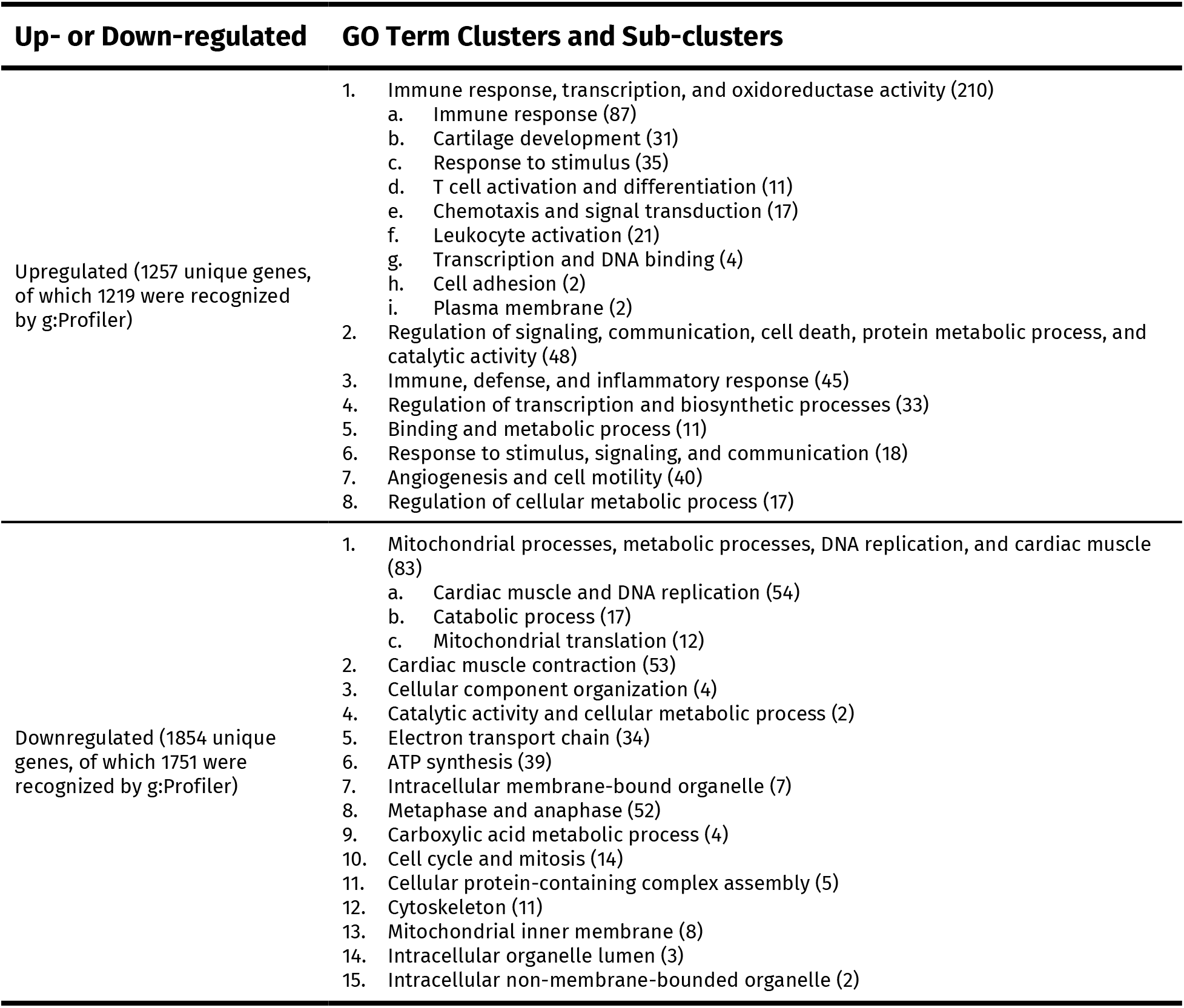
Summary of results of the hiPSC-CM cell line analysis. The names of each GO term cluster and sub-cluster from the hiPSC-CM cell line analysis (both upregulated and downregulated results) are displayed in the table. The number of GO terms that are in each cluster and sub-cluster is in parentheses. In the “Up- or Down-regulated” column, the number of uniquely regulated genes and the number of which were recognized by g:Profiler are also in parentheses.

Among these clusters, Cluster 1 (“Immune response, transcription, and oxidoreductase activity”) contained 210 GO terms and was therefore used to create sub-clusters. These GO terms formed 9 sub-clusters (Table 8, Figure S5, and Table S7B), 4 of which were unique to cardiomyocytes: “Cartilage development”, “Chemotaxis and signal transduction”, “Transcription and DNA binding”, and “Plasma membrane”. The remaining 5 GO clusters were: “Immune response”, “Response to stimulus”, “T cell activation and differentiation”, “Leukocyte activation”, and “Cell adhesion”.

1860 downregulated genes were identified from the hiPSC-CM cell line, of which 1854 were unique (Table S5C). These resulted in 321 total GO terms (211 BP terms, 85 CC terms, and 25 MF terms) (Table S3D) that could be grouped into 15 clusters (Table 8, Figure S6, and Table S8A). The 15 clusters of GO terms that represent the downregulated hiPSC-CM results were: “Mitochondrial processes, metabolic processes, DNA replication, and cardiac muscle”, “Cardiac muscle contraction”, “Cellular component organization”, “Catalytic activity and cellular metabolic process”, “Electron transport chain”, “ATP synthesis”, “Intracellular membrane-bound organelle”, “Metaphase and anaphase”, “Carboxylic acid metabolic process”, “Cell cycle and mitosis”, “Cellular protein-containing complex assembly”, “Cytoskeleton”, “Mitochondrial inner membrane”, “Intracellular organelle lumen”, and “Intracellular non-membrane-bounded organelle”.

Among the clusters of the downregulated hiPSC-CM results, Cluster 1 (“Mitochondrial processes, metabolic processes, DNA replication, and cardiac muscle”) contained 83 GO terms and was therefore split up into 3 sub-clusters (Table 8, Figure S7, and Table S8B): “Cardiac muscle and DNA replication”, “Catabolic process”, and “Mitochondrial translation”.

## Discussion

Since the beginning of the outbreak, SARS-CoV-2 has been heavily studied and researched in anticipation of understanding the disease and developing proper treatments. Generally, the most common symptoms of SARS-CoV-2 are fever, shortness of breath, cough, and loss of taste and smell, lasting 7-14 days on average [25]. Reports have shown that the virus is capable of not only affecting lung function, showing similar results as pneumonia, but also influences the function of various organs throughout the body, such as the heart, kidney, testis, liver, lymphocytes and nervous system [26], leading to a plethora of other symptoms of variable nature and severity in patients. Previous research has also shown that infection with SARS-CoV-2 leads to activation and upregulation of various aspects of the immune system, including recruiting cytokines and chemokines, as well as cell migration and degranulation [27]. Other studies have established the downregulation of several genes upon viral infection, such as angiopoietin-1, a powerful protector of the vascular system [28] and retinoic acid-induced protein 2, a gene responsible for cellular development and growth [24]. However, no studies have established the common set of genes that are differentially regulated by viral infection, and so, we have not been able to identify the specific effects of the virus on various cells and tissues in the body. To address this, we compared gene expression across a number of different cell types, and established the common and unique sets of genes regulated by viral infection.

We found the “core” transcriptome of SARS-CoV-2 infection to consist of 866 genes that were commonly upregulated across all tissue types (Table S2A). These genes mainly consisted of those involved in immune functions, including leukocyte activation, cytokine signaling and response to stimuli (Figure 5). Ion transporter activity also appeared to be commonly upregulated. Interestingly, only nine genes in total were commonly repressed by viral infection, spanning a range of functions such as inhibition of endothelial permeability, contractile activity, cellular growth, organization of cytoskeleton, and cell differentiation (Table 2). The low number of downregulated genes might be due to several different factors, one of which could be that the infected samples were all examined within 72 hours post SARS-CoV-2 infection, leaving little time for significant downregulation to be observed. The virus might mostly be upregulating the expression of genes within the initial time period after infection, leading to these skewed results. More work will need to be done to identify the causes for this discrepancy and to better establish the core set of commonly downregulated genes. We also found that viral infection did not abolish the inherent differences in gene expression profiles across cell types, since samples from a given cell type clustered together, irrespective of their viral infection status (Figure 3).

The results indicate that even though there is cell-type specific variance in the cellular responses of each individual cell line, there are common gene ontology clusters and pathways that can be utilized and targeted to inhibit the progression of SARS-CoV-2. This is especially true since SARS-CoV-2 is able to infect different body organs. The six clusters as well as the eight sub-clusters that were identified as commonly upregulated consist of GO terms that are related to immune system activation post-SARS-CoV-2 infection—more specifically, they were related to cytokine and chemokine activity (e.g. inflammatory response, immune cell activation, secretion, degranulation, and chemotaxis) (Table S4A and Table S4B). Past studies have shown a similar impaired upregulation of cytokines and chemokines in SARS-CoV-2 patient samples [29, 30].

We also found that ion transporter activities, especially potassium channels and calcium channels, were elevated across all cell types after SARS-CoV-2 infection. In cardiomyocytes, ion channel activities are critical in maintaining normal heart function by controlling heart contraction [31]. After infection, the expression of potassium and calcium channels is increased in cardiomyocytes, which could potentially have negative effects on heart functions, given that a correct cardiac rhythm needs precise control of ion conduction. It has also been shown that patients infected with SARS-CoV-2 exhibited various heart problems, including cardiac arrest, cardiomyopathy, and heart failure, which could be correlated with the abnormal ion channel activity [32].

Our results show how SARS-CoV-2 infection may possibly induce complications of existing diseases such as cancers [34, 35], or be exacerbated by existing conditions. For example, the universal downregulation of Myosin light chain kinase (MLCK), which plays a role in many different cellular processes, including cell migration, endocytosis, muscle contraction, and epithelial and endothelial barriers, may lead to various complications in patients [36].

Having established the “core” transcriptome, we could then analyze gene regulation in various cell types to identify genes that were uniquely up or downregulated upon viral infection. For example, in the kidney-derived HEK 293T cells (Table S6), we found that transcription and DNA binding activities, and cellular metabolism were upregulated specifically in these cells, allowing us to isolate the unique effects of the virus on these cells. Similarly, we found that regulation of transcription and biosynthetic processes, and binding and metabolic processes were upregulated in hiPSC-CM cells, and that functions like cardiac muscle contraction, cell cycle and mitosis, and mitochondrial activities were specifically downregulated in these cells (Table 8). Our results lay the foundation for better understanding the complicated interactions [37, 38] that may underlie the many organ system failures observed as one of the symptoms of COVID-19 [26, 39]. By establishing the cell-type specific changes to gene regulation, we help identify the likely mechanisms by which specific organ systems are damaged due to SARS-CoV-2 infection.

In lung-derived cells (NHBE), we observed a significant upregulation of MAF bZIP transcription factor F (MAFF—also upregulated in response to hypoxia), which has been shown to be fundamental in the activation of other genes involved in responses to cellular stress [40]. Interleukin 32 (IL32) which serves many roles, including inflammatory response to pathogens [41] was also uniquely upregulated in NHBE cells, indicating a role for this gene in the specific lung-related pathophysiology of COVID-19. In addition, 2′-5′-oligoadenylate synthetase 3 (OAS3) was also upregulated in these cells. This gene encodes the OAS3 enzyme of the 2′-5′-oligoadenylate synthetase family and plays a crucial role in inhibiting cellular synthesis during viral replication by degrading viral RNA [42, 43]. In multiple studies, NHBE cells have been shown to play a crucial role in initiating antiviral innate immune response, mediating host defense, and viral pathogenesis following infection by secreting cytokines and chemokines [44, 45]. Therefore, it is highly likely that the enrichment of MAFF, IL32, and OAS3 are attributed to cellular stress and innate immune response as a result of viral infection with SARS-CoV-2. More specifically, MAFF and OAS3 may play a major part in mediating host response, while IL32 is involved in hindering the virus’s pathogenic mechanism by apoptosis or other means.

Only three uniquely downregulated genes were identified in the NHBE cell line upon SARS-CoV-2 infection. Interferon induced transmembrane proteins (IFITM), including the downregulated IFITM10, restrain pathogenic entry into the host cells by suppressing viral membrane fusion and possibly hinder viral replication by preventing its gene expression and protein synthesis [46]. It is found within NHBE cells abundantly, and with significant downregulation of the gene, SARS-CoV-2 has unrestricted access to the cells, allowing for infection and consequential spreading [47]. Also found to be uniquely downregulated is C-X-C motif chemokine ligand 14 (CXCL14). CXCL14 is generally known to be an immune and inflammatory modulator, which performs various functions, suggesting that this gene engages the inflammatory response within NHBE cells [48, 49]. CXCL14 also significantly affects antimicrobial immunity [48]. Finally, MAP7 domain containing 2 (MAP7D2) is a microtubule-associated protein that is broadly known to stabilize microtubules and is broadly important for all microtubule-related functions [50]. Specifically, MAP7D2 is involved in the organization of the microtubule cytoskeleton, and downregulation of MAP7D2 could result in significant defects in axonal growth and diminished cargo entry [51]. Viral downregulation of this gene could result in increased infection of lung cells by the virus, possibly by breaking down cytoskeletal barriers to infection.

The main unique response of HEK 293T cells to SARS-CoV-2 infection is the upregulation of cell metabolism activity and cellular transcriptional activity (Table S6), suggesting that the regulation of these processes is particularly important to the effects of SARS-CoV-2 on these cells. Studies have also shown that the altered metabolism in kidney cells is closely associated with various kidney diseases [52], and our results imply that the kidney cells’ misregulation of metabolism in response to the virus could correlate with the kidney failure observed in some of the infected patients. In contrast to the 262 uniquely upregulated genes, there are only 12 uniquely downregulated genes in the HEK 293T cell line after SARS-CoV-2 infection. Some genes have not been well-documented, whereas the remainder play different roles and their downregulation may cause pathogenic effects on the kidney. For example, FPGT, PDK1, and MMAA encode proteins important in cell metabolism, and STEAP1 and LINC00472 are associated with cancer development (Table S5B) [53, 54]. While our results uncover some clues about the effects of SARS-CoV-2 on kidney cells, we must bear in mind that HEK 293T cells and kidney cells are not equivalent, and the actual responses of kidney cells to SARS-CoV-2 still need to be verified.

The uniquely upregulated genes from the hiPSC-CM cell line analysis and those from the combined analysis both share involvement in immune responses, indicating that different organ systems might elicit different responses, or activate similar pathways using different mechanisms. However, those from the hiPSC-CM cell line analysis were also involved in blood vessel development, metabolic and biosynthetic processes, transcription, and oxidoreductase activity. An upregulation in genes involved in angiogenesis in hiPSC-CMs aligns with a previous study that revealed cardiomyocytes as critical targets for SARS-CoV-2 infection [55]. Studies have shown that infection with SARS-CoV-2 damages blood vessels as the virus replicates within the hiPSC-CMs [56]. Therefore, the upregulation of blood vessel development in the hiPSC-CM cell line is consistent with the cardiovascular system trying to repair itself after viral infection.

It has been widely found that cardiomyocytes show significant ACE2 and spike protein expression following infection with SARS-CoV-2, which can lead to significant cardiac injury possibly associated with mortality [15]. It is suggested that the viral genome of SARS-CoV-2 is transcribed within cardiomyocytes following analysis of cardiac cells performed on autopsy specimens which showed significant presence of ACE2 receptors, as well as SARS-CoV-2 viral proteins within cardiac tissue [57]. Research shows that when viral information is transcribed within the hiPSC-CMs, its proteins are able to upregulate the cells’ transcription factors, as well as other regulatory genes, in turn disrupting the homeostasis of the hiPSC-CM and ultimately leading to apoptosis [58]. Our research shows that regulation of transcription, as well as cell death, is upregulated within the cell line. In cases where substantially large amounts of cell death are occuring, significant cardiac diseases, such as myocardial infarction could result [59].

Studies have shown that some COVID-19 patients exhibited cardiovascular abnormalities, such as acute cardiac injury and cardiogenic shock, but the relationship between SARS-CoV-2 infection and heart problems is still unclear [31, 60, 61]. While a lot of studies have been focusing on the systemic effects of the SARS-CoV-2 infection on cardiovascular functions, we focus on the intrinsic and unique cellular responses of cardiomyocytes to SARS-CoV-2 infection using hiPSC-CM cell line. Compared to the uniquely upregulated genes in hiPSC-CMs after infection, the downregulated results mostly feature the processes of mitochondrial activities, heart muscle contraction, and mitosis.

Mitochondria are crucial for cells to maintain normal activities. Studies have found that dysfunction of mitochondrial activities in cardiac and skeletal muscles are associated with heart failure [62]. Therefore, the downregulation of mitochondrial activity in cardiomyocytes after SARS-CoV-2 infection may indicate a similar cardiovascular response in the human body. Furthermore, we have also observed a downregulation in genes associated with heart muscle contraction. This is in accordance with a study which has found that hiPSC-CMs stop beating 72 hours after SARS-CoV-2 infection [15]. In the human body, the contraction of cardiomyocytes is critical in maintaining heart functions, and the abnormal heart contraction could lead to detrimental effects including arrhythmia, which is also a symptom of COVID-19 [31, 63].

Given that cell regeneration, specifically cytokinesis, rarely happens in mature human cardiomyocytes [64, 65], it is unclear how our observation that mitosis-associated genes are specifically downregulated in hiPSC-CMs relates to cardiomyocyte function *in vivo*. One explanation for this result could be the discrepancy in the nature between hiPSC-CMs and actual human cardiomyocytes. hiPSC-CMs can be induced to proliferate, and thus maintain the capability to undergo cell cycles and mitosis [66], whereas mature cardiomyocytes do not. However, our results might be relevant in the case of pregnant patients, where viral infection might have unpredicted effects on the fetus.

In addition to the lung-associated symptoms seen in COVID-19, a number of other organ systems are also affected by this disease. Several studies have noted non-respiratory COVID-19 symptoms such as kidney dysfunction, central nervous system tissue damage, elevated levels of particular enzymes that lead to liver damage, and arrhythmia [67, 68]. Moreover, multiple studies have established a number of common comorbidities that affect disease severity and outcomes, and these include obesity, hypertension, diabetes, and cardiovascular, cerebrovascular, respiratory, and kidney diseases [69, 70]. By utilizing our “core” transcriptome, we have established tissue-specific gene dysregulations that shed some light on the mechanisms by which SARS-CoV-2 might cause damage to different organs, or interact with existing chronic conditions. Current therapeutics recommended by the NIH include dexamethasone, which has been found to reduce mortality in patients who require supplemental oxygen, with the greatest effect observed in those who require mechanical ventilation, and can be administered in combination with remdesivir, the only treatment for COVID-19 approved by the FDA [71]. However, current treatment regimes lack recommendations to address non-respiratory symptoms. Since our study analyzed a variety of patient samples and cell types, including cells found in the respiratory system, our results can point the way for researchers to study and design therapeutics that relieve both respiratory and non-respiratory symptoms in hospitalized COVID-19 patients.

Our strategy of using RNA-seq to identify upregulated and downregulated gene ontologies and genes from a variety of cell types can be applied to other coronavirus diseases. Considering that SARS-CoV, MERS-CoV, and SARS-CoV-2 are all coronaviruses that primarily target the respiratory system, most preliminary research is focused on how the respiratory system is affected in patients, leaving gaps in non-respiratory research. Comprehensive studies similar to ours can assist in filling the gaps in research by providing clues as to which respiratory and non-respiratory functions are being upregulated and downregulated as a result of infection. This can help to address non-respiratory severe symptoms such as kidney failure, which was a significant comorbidity for SARS, MERS, and COVID-19 [72]. Our findings drive home the conclusion that the more we learn about the human body’s response to SARS-CoV-2 infection and the symptoms it may cause, the better position we all will be in to thwart the coronavirus and any others that might emerge in the future.

