## Supplemental information for "Identifying The “Core” Transcriptome of SARS-CoV-2 Infected Cells"

### Supplemental Materials

**NOTE:** We have included links to spreadsheets for all lists in the tables for the sake of readability, rather than have some tables with information on >1,000 genes.

**Table S1.** Table of BioProjects and SRRs used in the experiment. Lists the BioProjects and RNA Seq information used in this experiment.

**(A)**

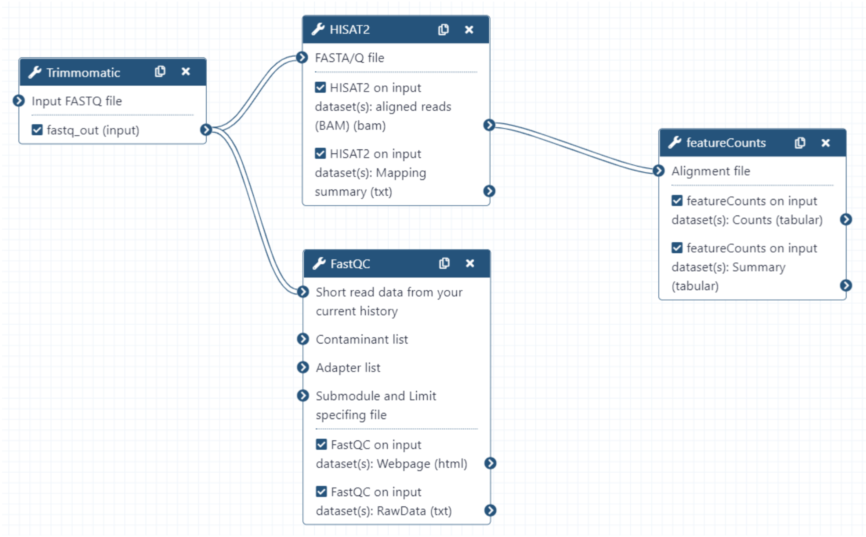


**(B)**

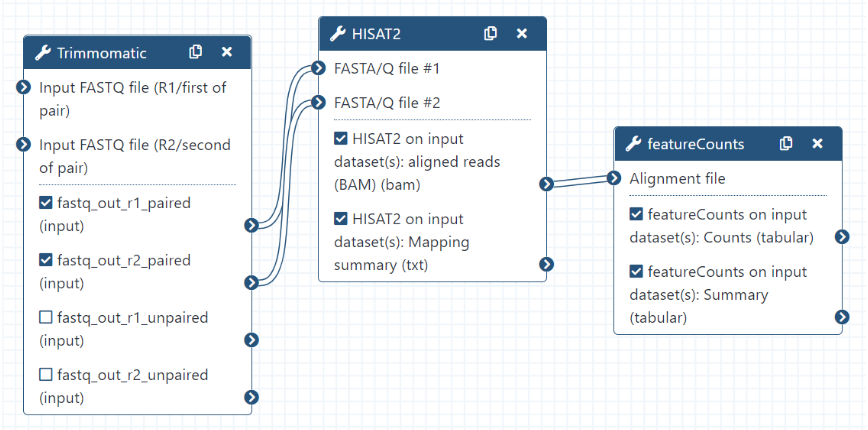


**Figure S1. Galaxy workflow for (A) single-end and (B) paired-end SRR datasets.** The raw RNA Seq data was run through Galaxy workflow (A) for single-end or (B) paired-end RNA Seq, to obtain differentially regulated genes. These workflows were used in both two-factor analysis and one-factor analysis.

**Table S2. List of (A) upregulated genes and (B) downregulated genes from the combined analysis.** (A) 866 genes are differentially upregulated and (B) 9 genes are differentially downregulated across all or most of the cell lines and patient samples. The genes were obtained by performing the DESeq2 in Galaxy ($P_{adj} < 0.01$; $Log_2FC < -1$ or $Log_2FC > 1$). The gene names and descriptions are obtained from gProfiler.

**Table S3. Binary matrices for (A) Combined upregulated results and cluster 1, (B) HEK 293T upregulated results, (C) hiPSC-CM upregulated results and cluster 1, and (D) hiPSC-CM downregulated results and cluster 1.**

**Table S4. (A) Spreadsheet of GO term clusters obtained from combined analysis (upregulated genes) and (B) GO term sub-clusters obtained from combined analysis (upregulated genes) cluster 1 ("Immune, defense, and inflammatory response").** Table of clusters deduced using R program based on molecular function, biological process, and cellular component. Each cluster has its own color and representative GO term. (A) Jaccard clustering was used for the upregulated combined experiment's GO analysis, resulting in 6 clusters. (B) Cluster 1 ("Immune, defense, and inflammatory response") was further analyzed using eDice clustering, resulting in 8 clusters.

**Table S5. List of unique genes for (A) NHBE, (B) HEK 293T, and (C) hiPSC-CM (gene IDs, names, GO functions).** Tables of upregulated and downregulated genes that are unique to either the (A) NHBE cell line, (B) HEK 293T cell line, or (C) hiPSC-CM cell line. The lists of differentially regulated genes from the NHBE, HEK 293T, and hiPSC-CM cell line analyses, separately, were compared with the list of differentially regulated genes from the combined experiment in order to determine which genes were uniquely differentially expressed by the NHBE, HEK 293T, and hiPSC-CM cell lines. Each table lists each gene with its corresponding ID, name, and description. A short statement or description of their functions can be seen in the inset found in the cell of the gene ID.

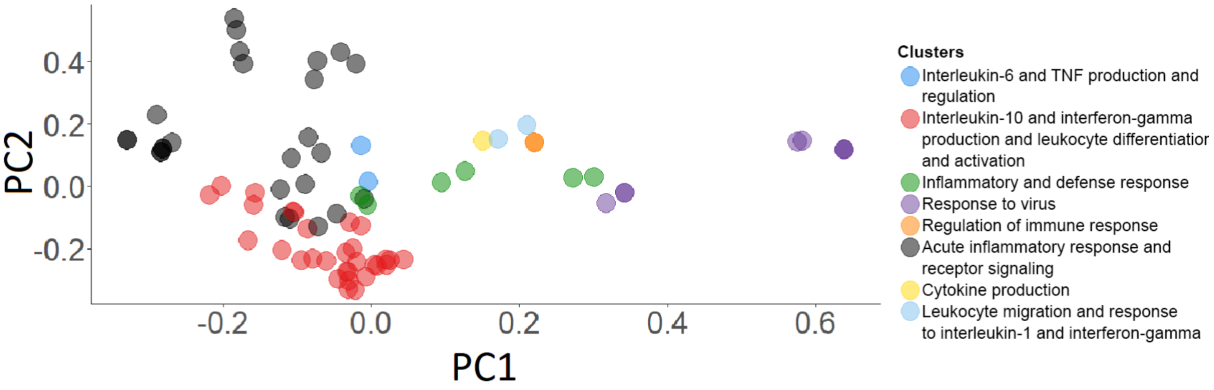


**Figure S2. Principal Component Analysis (PCA) plot for 77 GO terms ($P_{adj} < 0.01$) from Cluster 1 of the combined analysis.** They are further categorized into eight sub-clusters, represented with different colors, using the R code with the eDice algorithm.

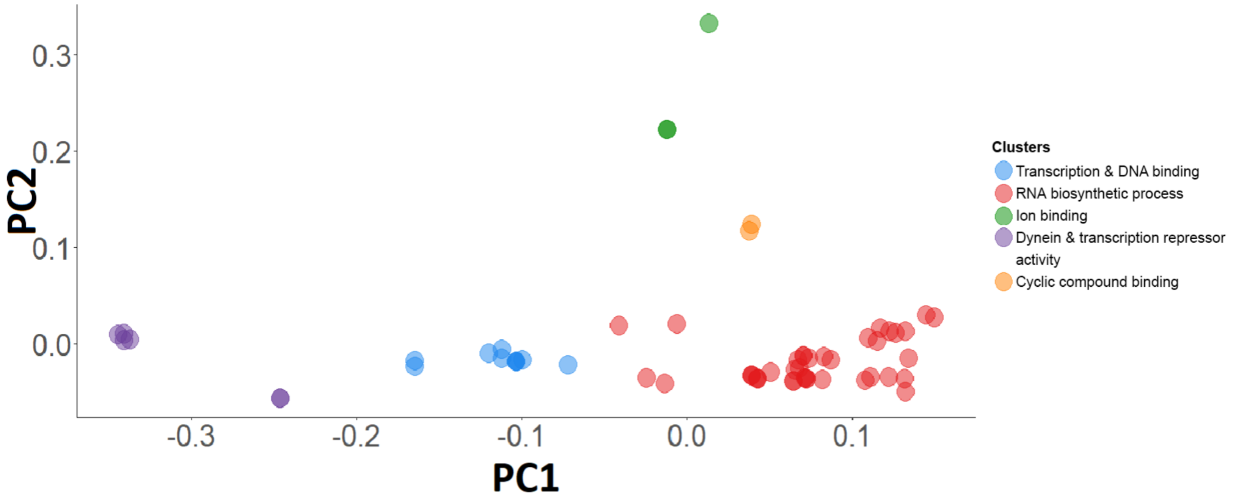


**Figure S3. Principal Component Analysis (PCA) plot for 62 GO terms ($P_{adj} < 0.01$) of uniquely upregulated genes in the HEK 293T cell line.** They are categorized into 5 clusters, represented with different colors, using the R code with Gower algorithm.

**Table S6. Spreadsheet of GO term clusters obtained from HEK 293T cell line analysis (upregulated genes).** Table of clusters deduced using R program, Gower clustering, based on molecular function, biological process, and cellular component of uniquely upregulated genes for the HEK 293T cell line, resulting in 5 clusters. Each cluster has its own color and representative GO term.

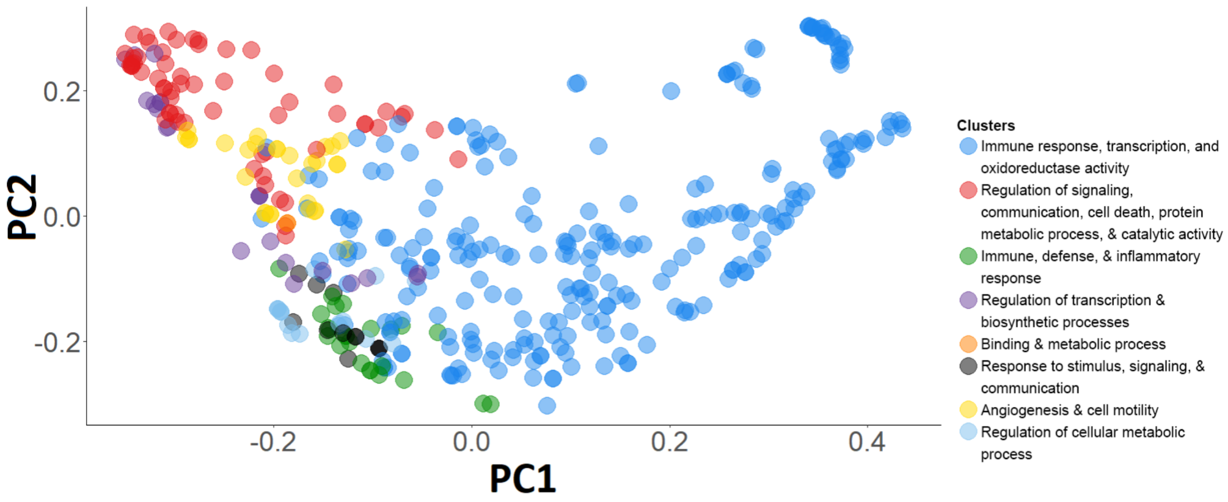


**Figure S4. Principal Component Analysis (PCA) plot for 422 GO terms ($P_{adj} < 0.01$) of uniquely upregulated genes in the hiPSC-CM cell line.** They are categorized into 8 clusters, represented with different colors, using the R code with Jaccard algorithm**.**

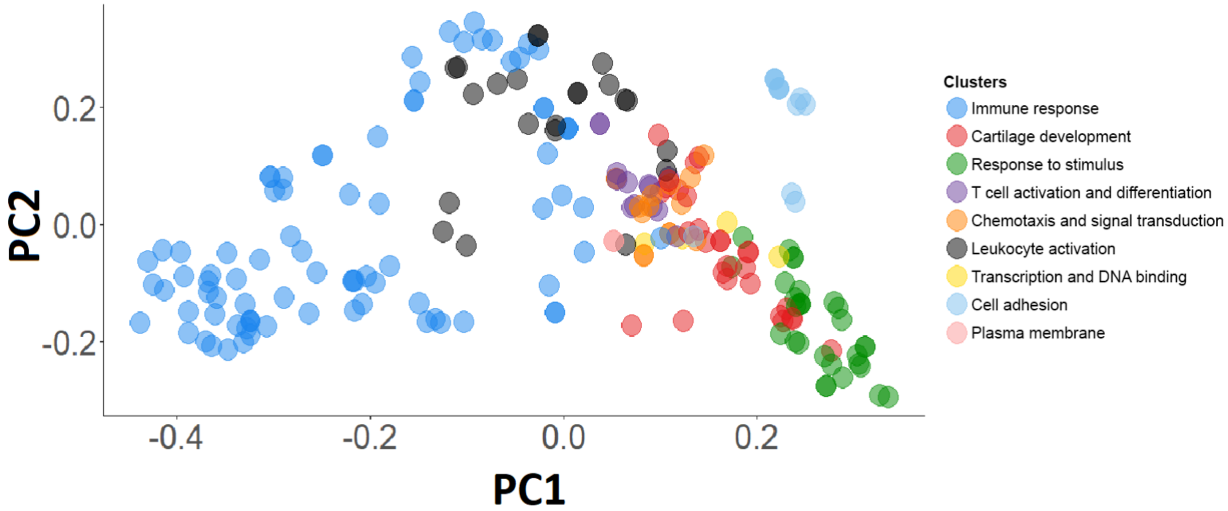


**Figure S5. Principal Component Analysis (PCA) plot for 210 GO terms ($P_{adj}$ < 0.01) from Cluster 1 of hiPSC-CM upregulated analysis.** They are further categorized into 9 sub-clusters, represented with different colors, using the R code with the Jaccard algorithm.

**Table S7. (A) Spreadsheet of GO term clusters obtained from hiPSC-CM cell line analysis (upregulated genes) and (B) GO term sub-clusters obtained from hiPSC-CM cell line analysis (upregulated genes) cluster 1 ("Immune response, transcription, and oxidoreductase activity").** Table of clusters deduced using R program based on molecular function, biological process, and cellular component. Each cluster has its own color and representative GO term. (A) Jaccard clustering was used for the upregulated hiPSC-CMs cell line GO analysis, resulting in 8 clusters. (B) Cluster 1 ("Immune response, transcription, and oxidoreductase activity") was further analyzed using Jaccard clustering, resulting in 9 clusters.

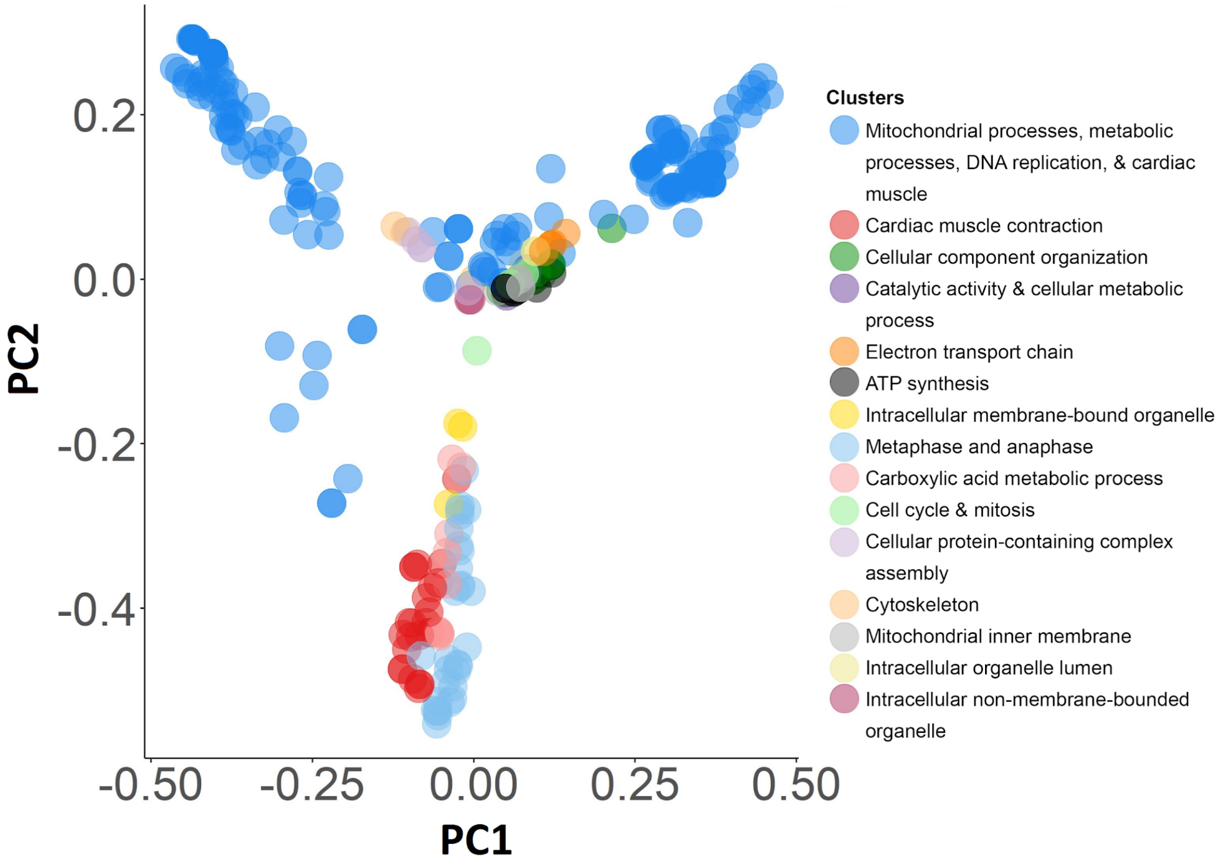


**Figure S6. Principal Component Analysis (PCA) plot for 321 GO terms ($P_{adj} < 0.01$) of uniquely downregulated genes in the hiPSC-CM cell line.** They are categorized into 15 clusters, represented with different colors, using the R code with the eDice algorithm.

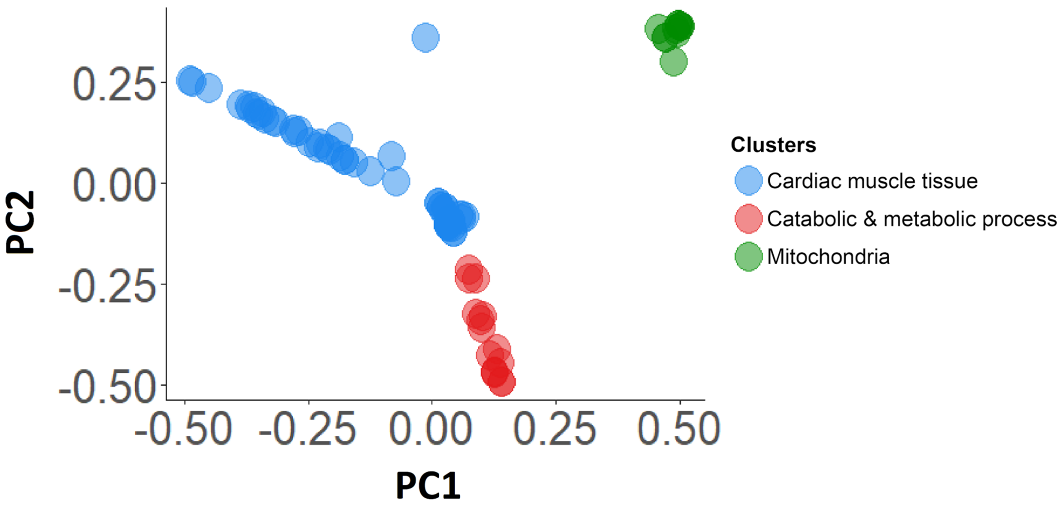


**Figure S7. Principal Component Analysis (PCA) plot for 83 GO terms from Cluster 1 of hiPSC-CM downregulated analysis.** They are further categorized into 3 sub-clusters, represented with different colors, using the R code with the eDice algorithm.

**Table S8. (A) Spreadsheet of GO term clusters obtained from hiPSC-CM cell line analysis (downregulated genes) and (B) GO term sub-clusters obtained from hiPSC-CM cell line analysis (downregulated genes) cluster 1 ("Mitochondrial processes, metabolic processes, DNA replication, and cardiac muscle").** Table of clusters deduced using R program based on molecular function, biological process, and cellular component. Each cluster has its own color and representative GO term. (A) eDice clustering was used for the downregulated hiPSC-CMs cell line GO analysis, resulting in 15 clusters. (B) Cluster 1 ("Mitochondrial processes, metabolic processes, DNA replication, and cardiac muscle") was further analyzed using eDice clustering, resulting in 3 clusters.
